## Supplemental Information for "Differential PaxillinB dynamics at *Dictyostelium* cell-substrate adhesions"

**SUPPLEMENTARY INFORMATION**

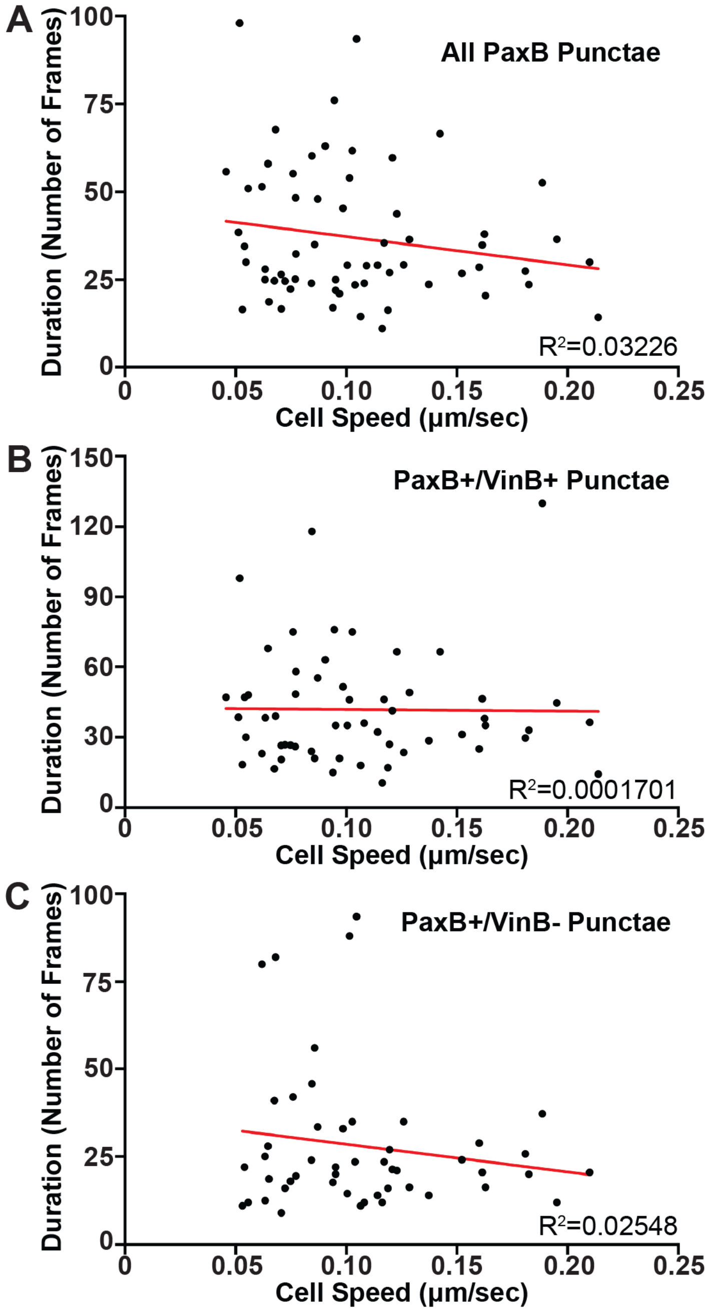


**Supplemental Figure 1: *Dictyostelium* cell-substrate adhesion duration does not correlate with cell migration speed**

Scatterplot graphs comparing cell migration speed to the duration of **A)** PaxillinB-positive punctae, **B)** PaxillinB+/VinculinB+ punctae or **C)** PaxillinB+/VinculinB- punctae across n = 65 (A), 57 (B), and 48 (C) cells, respectively. Correlation analyses of cell migration speed and punctae duration suggest there is no correlation between punctae duration and cell migration speed regardless of adhesion composition.
